# yEvo: Experimental evolution in high school classrooms selects for novel mutations and epistatic interactions that impact clotrimazole resistance in *S. cerevisiae*

**DOI:** 10.1101/2021.05.02.442375

**Authors:** M. Bryce Taylor, Ryan Skophammer, Alexa R. Warwick, Josephine M. Boyer, yEvo Students, Margaux Walson, Christopher R. L. Large, Angela Shang-Mei Hickey, Paul A. Rowley, Maitreya J. Dunham

## Abstract

Antifungal resistance in pathogenic fungi is a growing global health concern. Non-pathogenic laboratory strains of *Saccharomyces cerevisiae* are a useful model for studying mechanisms of antifungal resistance that are relevant to understanding the same processes in pathogenic fungi. We developed a series of lab modules in which high school students used experimental evolution to study antifungal resistance by isolating azole-resistant *S. cerevisiae* and examining the genetic basis of resistance. All 99 sequenced clones from these experiments possessed mutations previously shown to impact azole resistance, demonstrating the efficacy of our protocols. We additionally found recurrent mutations in an mRNA degradation pathway and an uncharacterized mitochondrial protein (Csf1) that have possible mechanistic connections to azole resistance. The scale of replication in this high school-led initiative allowed us to identify epistatic interactions, as evidenced by pairs of mutations that occur in the same clone more frequently than expected by chance (positive epistasis) or less frequently (negative epistasis). We validated one of these pairs, a negative epistatic interaction between gain-of-function mutations in the multidrug resistance transcription factors Pdr1 and Pdr3. This high school-university collaboration can serve as a model for involving members of the broader public in the scientific process to make meaningful discoveries in biomedical research.

## Introduction

Azoles are the primary class of antifungals used in medicine and agriculture. Azole resistance in fungi is a growing global health crisis (Fisher *et al.* 2018). Characterizing the genetic basis of azole resistance provides an opportunity to predict whether a newly-observed clinical isolate will have resistance to commonly-utilized antifungals and may reveal candidates for new treatment paradigms (Usher and Haynes 2019; Cowen *et al.* 2009; Song *et al.* 2020).

Like many antifungal drugs, azoles target a component of cell membrane biogenesis. Azoles competitively bind the active site of the enzyme Erg11, preventing a key rate-limiting step in sterol biosynthesis (Veen *et al.* 2003). Ergosterol in yeasts is functionally equivalent to human cholesterol and the production pathway is highly conserved between humans and yeast (Kachroo *et al.* 2015 Science). Like cholesterol, ergosterol is a key component of the cell membrane, influencing its fluidity, permeability, and organization (Dufourc 2008; Hannich *et al.* 2011). Depletion of membrane ergosterol by inhibiting Erg11 is fungistatic (prevents growth) and leads to the accumulation of toxic intermediates of sterol biosynthesis (Kelly *et al.* 1995; Allen *et al.* 2015).

Due to the widespread use of azoles as therapeutics and prophylactics, azole drug resistance has been studied in detail in various species of fungi for decades. Azole resistance mutations fall into two overarching classes. The first class compensates for Erg11 inhibition through the increased production of the Erg11 enzyme (to overcome the competitive inhibition by azoles) or by alterations to the Erg11 active site (to prevent enzyme inhibition by azoles). This can be accomplished through *ERG11* gene duplication, altered activity of regulators of *ERG11* expression, or point mutations in the enzyme itself (Berkow and Lockhart 2017). The second class of mutations, referred to as pleiotropic drug or multidrug resistance mutations (Balzi and Goffeau 1991; Gulshan and Moye-Rowley 2007), lead to increased production or activity of efflux pumps such as Pdr5 (Wolfger *et al.* 2001; Kumari *et al.* 2021). This can be accomplished through point mutations in the pleiotropic drug response transcription factors Pdr1 and Pdr3, loss of repressors of *PDR5* expression, or point mutations in the pump itself (Balzi and Goffeau 1991; Gulshan and Moye-Rowley 2007).

Though the budding yeast *S. cerevisiae* is only an opportunistic pathogen under rare circumstances (Clemons *et al.* 1994), it shares drug resistance pathways with pathogenic fungi and has been a useful model for clarifying drug resistance mechanisms beyond the major mutations described above (Paul and Moye-Rowley 2014; Demuyser and Van Dijck 2019). For instance, the activity of Pdr transcription factors, particularly Pdr3 in *S. cerevisiae* (Hallstrom and Moye-Rowley 2000; Zhang and Moye-Rowley 2001), is repressed by respiration; mutations that shift metabolism toward fermentation, such as petite mutations that result in a respiratory deficiency, can relieve this repression and promote *PDR5* expression and drug resistance. Also, ergosterol production is regulated by several pathways including oxygen sensing (Serratore *et al.* 2018) and the RAM (Regulation of Ace2 and Morphogenesis) network which regulates cell division and is conserved from yeast to humans (Nelson *et al.* 2003; Walton *et al.* 2006; Mulhern *et al.* 2006; Saputo *et al.* 2012). Mutations in components of these pathways can increase resistance to azoles by increasing production of ergosterol, which interacts with sphingolipids in the cell membrane. Alterations to sphingolipid concentration can allow a cell to better tolerate azole exposure (Francois *et al.* 2009).

The majority of genetic studies of azole resistance have utilized traditional mutant selection or clinical isolate screening approaches that tend to focus on single, strong-effect mutations. Experimental evolution provides an opportunity for the identification of mutations with a small or background-dependent effect. Previous evolution experiments selecting for azole resistance have demonstrated that this paradigm can identify new resistance factors and can validate candidate secondary antifungals that prevent resistance evolution (e.g. Cowen *et al.* 2000, Anderson *et al.* 2003, Selmecki *et al.* 2009, Hill *et al.* 2013, Boyer *et al.* 2021). We anticipated that additional replicates carried out with different culturing protocols would lead to the identification of novel resistance factors.

The wealth of information available on azole resistance mechanisms and possibility of identifying new resistance factors makes this system particularly attractive for a course-based research module. To leverage this, we developed protocols suitable for carrying out experimental evolution of azole resistance in high school classrooms as part of a course-based research experience called yEvo (available at yevo.org). We developed partnerships with teachers at two high schools to ensure these protocols were compatible with their classrooms and learning objectives. The laboratory activities explicitly connect evolution to underlying molecular biology in an open-ended inquiry framework and provide a powerful demonstration of how evolution occurs at the molecular level. Connecting these topics is a key goal of the Next Generation Science Standards (NGSS Lead States 2013), a current benchmark for K-12 science education in the United States.

Our pedagogical aims and evaluation will be reported in detail elsewhere (Taylor *et al.* in preparation). In this paper, we characterize 99 evolved clones from these experiments using phenotyping assays and whole genome sequencing. We were able to isolate clotrimazole-resistant strains as early as 2 weeks into the evolution protocol, though the majority of experiments were continued for an entire school year (30 to 34 weeks), which allowed time for multiple mutations to arise in most clones. Evolved clones were enriched for mutations impacting known azole resistance factors, as well as in genes that had not previously been associated with azole resistance. We provide evidence that these do impact azole resistance, expanding our understanding of the genetics of this important trait. We further show that epistatic interactions between evolved mutations can impact the evolution of resistance.

## Materials and Methods

### Yeast strains

Evolution experiments were performed with lab-derived strains of *Saccharomyces cerevisiae*; Haploid replicates utilized the *MATa* S288C derivative BY4741 and the *MATɑ* S288C derivative BY4742. 5 replicates used a diploid S288c derived by mating BY4741 and BY4742. All strain genotypes are listed in (**Table S1**). Strains used in evolution experiments carried a 2μ plasmid with KanMX, which provides resistance to the general antibiotic G418, and a pigment production pathway that gives each strain a unique color (**Table S2**; courtesy of the Boeke lab at New York University). G418 was added to the media to select for maintenance of these plasmids and reduce risk of contamination.

#### Classroom protocols

##### Westridge School evolution protocol

Evolution experiments were carried out via batch transfer. Every 2-4 days yeast were transferred on a sterile cotton swab from a saturated culture to a tube of fresh media containing a concentration of clotrimazole of their choosing. Clotrimazole was obtained from an over-the-counter 1% solution dissolved in 70% isopropanol. Clotrimazole was added directly to culture tubes in the desired concentration prior to the introduction of yeast. Cultures were maintained in 5ml of YPD + G418 + clotrimazole at room temperature without aeration. Student groups maintained three replicates of an assigned strain throughout their AP Biology course. Yeast were initially exposed to a low dose of clotrimazole (2.25μM), which slowed but did not prevent ancestor growth, and doses were increased at 2x intervals (e.g. 4.5μM, 9μM, 18μM, etc.).

##### Moscow High School evolution protocol

Evolution experiments were carried out via batch transfer. Every 7 days yeast were transferred on a sterile cotton swab from a saturated culture to a tube of fresh media containing a concentration of clotrimazole of their choosing. Clotrimazole was obtained from an over-the-counter 1% solution dissolved in 70% isopropanol. Cultures were maintained in 5ml of YPD + G418 + clotrimazole at room temperature without aeration. Student groups maintained one replicate of an assigned strain throughout their Biology course. Yeast were initially exposed to a much higher dose (10μM), which prevented visible growth of the ancestor over short time scales (**Figure 1C**), but permitted growth over the extended time between transfers, and doses were increased at 2x intervals (e.g. 20μM, 40μM, 80μM, etc.).

**Figure 1.**
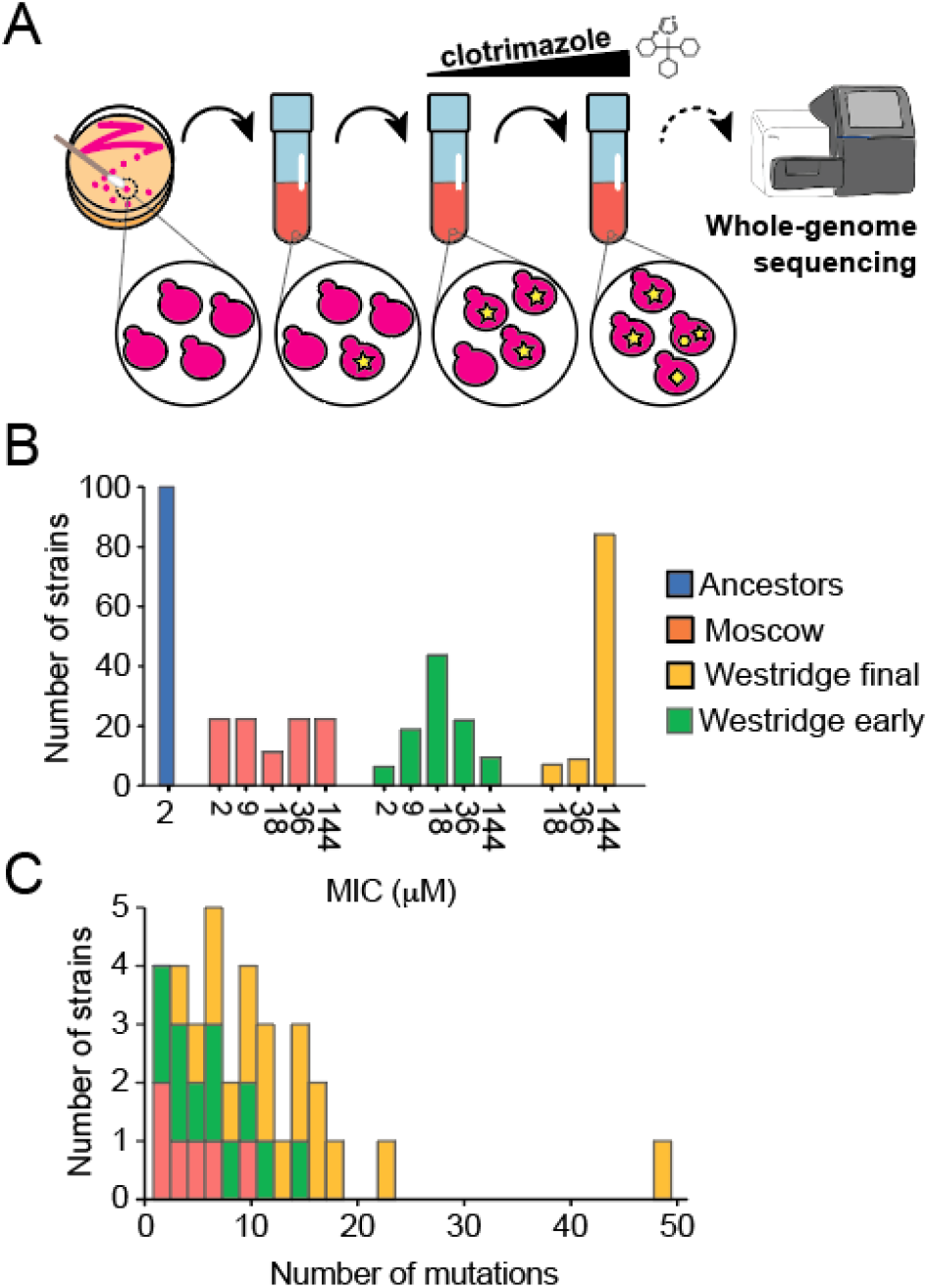
Overview of evolution experiment and sequencing. (A) outline of experiment. Yeast were propagated in increasing concentrations of clotrimazole for several weeks. Clones from these experiments were sequenced to identify mutations that occurred during the experiment. (B) Maximum measured clotrimazole tolerance of clones isolated from experiments at different schools and timepoints. (D) Number of point mutations in sequenced clones.

##### Strain storage

Freezer stocks of student populations at Westridge School were generated weekly by adding a 1:1 mixture of 50% glycerol solution in a cryogenic vial and stored in a non-frost-free −20°C freezer until transfer to −80°C at the University of Washington. Populations from Moscow High School were stored at 4°C until the end of the evolution protocol. Clones were selected and frozen at −80°C at the University of Idaho.

#### University lab protocols

##### Phenotyping of evolved clones

Clones from Westridge School were isolated from each Westridge School experiment at two timepoints (early and final, approximately 7 weeks and 30-34 weeks, respectively) and at weekly intervals from Moscow High School experiments. One clone was chosen from each student group for whole-genome sequencing. These were assayed for resistance to clotrimazole by minimum inhibitory concentration assay as follows: clones were grown for 48 hours at 30°C in 200ul of YPD + G418 medium in a 96-well plate without aeration. Cultures were resuspended, and 2ul of each culture was transferred to a new 96-well plate containing YPD + G418 medium with 9μM clotrimazole and monitored for growth over 2 days. To assay petite status, clones were additionally transferred to YPG (glycerol as carbon source) + G418 medium and monitored for growth over 3 days.

##### Whole-genome sequencing

Whole genome sequencing libraries were generated using a modified version of the Illumina Nextera protocol based on (Baym *et al.* 2015). Briefly, genomic DNA is fragmented using an Illumina Tagmentation enzyme that adds Illumina adaptor sequences to ends (Tagmentation). Tagmented samples are then PCR amplified using oligonucleotide primers that add unique barcodes to each sample so that samples can be multiplexed on a single Illumina sequencing run. Coverage of all clones described can be found in **Table S5**. Sequencing reads will be deposited in the NCBI Sequence Read Archive (SRA) upon publication and are available upon request.

##### Mutation calling

Briefly, reads were aligned to an S288c reference genome using BWA and mutations were called with Samtools. Mutations within 200 bases of a start codon were listed as putative promoter mutation. Single nucleotide polymorphisms with a quality score lower than 100 (out of 228) were discarded. All mutations with quality score less than 200, as well as all indels, were manually inspected in the Integrative Genomics Viewer (IGV) to ensure variants were present in a majority of reads from each mutant and were not present in reads from the respective ancestor. Common mutation targets were manually inspected in IGV to look for evidence of mutations that were missed by our mutation calling pipeline. We identified one such instance in the gene *PDR1* in strain: Westridge_C_final_1_2017-2018.

##### Copy number variants

Average coverage genome-wide was determined in 1000bp windows using the program IGVtools and plotted in R. Continuous regions with coverage that exceeded the genome-wide average by >= 2x in haploids were considered copy number variants. Regions that deviated from the genome-wide average by 0.5x in diploids were considered copy number variants.

##### Transposable element calling

Clone sequencing data were examined for evidence of mobilization events of transposable elements versus the reference genome using the McClintock pipeline (Nelson *et al.* 2017). Afterwards, comparisons were made between the ancestral strain and the clones to find exclusively *de novo* events within our experiments that were at least 1000-bp away from existing events with the software Bedtools. Subsequently, the transposition events were manually inspected in IGV for veracity using the split and discordant reads, generated using BWA and later processed with SAMblaster (Faust and Hall 2014) and samtools. One mobilization in the gene *PDR3* was confirmed by PCR and Sanger sequencing using primers PDR3_genotype_F and PDR3_genotype_R (**Table S3**).

##### Lineage determination

Clones from the same group that shared at least one point mutation were considered as coming from the same lineage. Shared mutations are denoted as non-independent in **(Table S6)**. Mutations that were shared by multiple members of a lineage were considered as single mutations events for the purpose of identifying recurrently mutated genes.

##### CRISPR/Cas9 allele replacements

A PAM (polyspacer adjacent motif) near *CSF1*^A2913P^ was targeted for cutting by the Cas9 enzyme. Oligonucleotides for guide RNA design and repair donors can be found in **Table S3**. Guide RNA oligonucleotides were introduced into pML104 backbone by Gibson assembly (**Table S2**). A G to C mutation was introduced to recreate the *CSF1*^A2913P^ allele detected in one evolved clone. To prevent recutting, a synonymous mutation that altered the PAM was introduced in codon A2913 on its own (control) or in combination with the A2913P mutation. Stationary yeast culture of *MATɑ* haploid S288c (**Table S1**) was transformed with 100ng Cas9 + gRNA expression vector and 1ug donor DNA with the lithium acetate protocol. Genotype of transformed clones at *CSF1* was determined by Sanger sequencing with primers CSF1_genotype_F + CSF1_genotype_R (**Table S3**).

##### CSF1 *allele replacement competitions*

Stationary phase cultures of wild type, synonymous mutant, and nonsynonymous mutant were mixed in equal proportions. 50ul of this mixture was inoculated into 5ml of either YPD or YPD + 9μM clotrimazole. These cultures were grown at 30°C in a roller drum until they reached stationary phase, approximately 2 days. 50ul of stationary culture was transferred to 5ml of respective media. This backdiluting was performed twice for a total of 3 outgrowths. Frequency of *CSF1*^A2913P^ was determined at initial and final timepoints by Sanger sequencing. Frequency of *CSF1*^A2913P^ allele at each sequenced timepoint was determined with the program QSVAnalyzer (Carr *et al.* 2009). Frequencies depicted are averages of 3 replicates. Error bars are one standard deviation in each direction (**Table S11**).

##### Tetrad dissections

Diploids were grown overnight in 5ml YPD in a roller drum at 30°C. 1ml of stationary-phase culture was pelleted and resuspended in 5ml sporulation medium as in (CSHL). These sporulation cultures were grown on a roller drum at room temperature. After 3-5 days, 50ul culture was pelleted and resuspended in 15ul YLE for 22 minutes @ 30°C. 500ul of water was slowly added to this mixture to dilute cells. 50ul of digested spore mixture was dripped down a YPD agar plate and allowed to dry. Tetrads were dissected with a micromanipulator microscope.

##### Restriction enzyme genotyping

Restriction fragment length polymorphisms were identified through manual examination of mutant and wild type sequences in SnapGene. Polymorphisms were amplified with Phusion polymerase with kit protocol. Primer sequences can be found in (Table S3). 8.5ul of PCR product was mixed with 1ul 10x cutsmart buffer + 0.25ul enzyme + 0.25ul water and digested for 1hr at 37C. Restriction enzymes RsaI and MboI were sourced from New England Biolabs.

##### Plate reader experiments

Genotyped spores were grown in 200ul YPD medium in 96-well plates for 48 hours. Cultures were resuspended and 2ul of each culture was transferred to 198ul YPD + 9μM clotrimazole media. Growth of these cultures was monitored in a plate reader for 24 hours at 30°C with orbital shaking. Average growth rate of all strains with the same genotype was calculated and plotted in R. Average maximum growth rate for each genotype was calculated with R library *growthrates*.

## Results

### Isolation of clotrimazole-resistant clones from a course-based research module

We implemented an experimental evolution lab module at two high schools in the United States. Protocols for these experiments were developed in close collaboration with each teacher to best match time, resources, and learning objectives (**Figure 1A**) (Taylor *et al.* in preparation). These experiments differed in key ways (**Methods; Figure 1B**). For instance, students at Westridge School carried out experiments throughout the school year with serial passage occuring every class period, or roughly every 2-4 days. Five replicates from Westridge School utilized diploid strains instead of the haploids utilized in all other experiments. Students at Moscow High School carried out experiments for an average of 11 passages at weekly intervals. At the conclusion of these experiments, evolved yeast populations were collected by partnering research laboratories at the University of Washington and the University of Idaho (**Methods**). Clotrimazole-resistant clones were isolated from an early time point (7 weeks at Westridge School and 2-12 weeks at Moscow High School), as well as from a later time point (30 to 34 weeks) at Westridge School. Clones were capable of growing in higher doses of clotrimazole than the unevolved ancestors. Clones from later time points were able to grow in high concentrations of azole with 53 of 57 late time point strains able to tolerate 144μM clotrimazole, compared to 14 of 42 early time point strains (**Figure 1C**; **Table S4**).

### Whole genome sequencing of evolved strains identified de novo mutations connected to azole resistance

The genome sequence of each confirmed drug-resistant isolate was determined using short-read next-generation sequencing. The genome sequences of the azole-resistant clones and their ancestors were aligned to the *S. cerevisiae* reference genome to identify sequence differences unique to evolved clones (**Methods**). We looked for point mutations (**Table S6**; **Table 1**), loss of mitochondrial DNA (ρ^0^ petite mutations) (**Table S8**), copy number changes (**Table S9**; **Figure 2**), and transposable element mobilizations (**Table S10**).

**Table 1.** Recurrently mutated genes. Genes with at least 3 independent nonsynonymous, indel, or nonsense mutations. Literature support is defined by at least one instance in which a gene or pathway has been implicated in resistance or sensitivity to an azole drug or in ergosterol production in a published research article.

| Gene | Unique events | # clones w/ mutation | Function | Literature support for gene? | Literature support for pathway? |
| --- | --- | --- | --- | --- | --- |
| *PDR1* | 33 | 65 | *PDR5* regulation | yes (Gulshan and Moye-Rowley 2007; Paul and Moye-Rowley 2014) | yes (Gulshan and Moye-Rowley 2007; Kumari *et al.* 2021; Paul and Moye-Rowley 2014) |
| *SUR1* | 30 | 44 | Sphingolipid production | yes (François *et al.* 2009; Vandebosc h *et al.* 2013; Kapitzky *et al.* 2010; Hoepfner *et al.* 2014) | yes (François *et al.* 2009; Gao *et al.* 2018) |
| *HAP1* | 26 | 30 | Oxygen sensing and ergosterol regulation | yes (Serratore *et al.* 2018) | yes (Jordá and Puig 2020) |
| *ERG25* | 25 | 49 | Ergosterol production | yes (Gachotte *et al.* 1997; Smith *et al.* 2016) | yes (Gachotte *et al.* 1997; Joseph-Horne and Hollomon 2006) |
| *PDR3* | 15 | 26 | *PDR5* regulation | yes (Gulshan and Moye-Rowley 2007; Paul and Moye-Rowley 2014) | yes (Gulshan and Moye-Rowley 2007; Kumari *et al.* 2021; Paul and Moye-Rowley 2014) |
| *CSG2* | 13 | 17 | Sphingolipid production | yes (Vandebosc h *et al.* 2013; Kapitzky *et* *al.* 2010;<br>Hoepfner *et al.* 2014) | yes (François *et al.* 2009; Gao *et al.* 2018) |
| *DHH1* | 6 | 8 | mRNA degradation | yes<br>(Vandenbosch *et al.* 2013) | no |
| *ROX1* | 6 | 6 | Oxygen sensing and ergosterol regulation | yes (Henry *et al.* 2002) | yes (Jordá and Puig 2020) |
| *TAO3* | 4 | 6 | RAM signaling | no | yes (Nelson *et al.* 2003;<br>Song *et al.* 2008; Saputo *et al.* 2012) |
| *ATP1* | 4 | 7 | Mitochondrial ATP synthesis | yes (Li *et al.* 2017) | yes (Li *et al.* 2017) |
| *ATP2* | 4 | 4 | Mitochondrial ATP synthesis | no | yes (Li *et al.* 2017) |
| *CSF1* | 4 | 5 | Unknown | yes (Mount 2018) | no |
| *DCP2* | 4 | 5 | mRNA degradation | no | no |
| *SIT4* | 4 | 5 | Regulates multidrug resistance and sphingolipid synthesis* | yes (Miranda *et al.* 2010;<br>Vandenbosch *et al.* 2013;<br>Khandelwal *et al.* 2018) | yes (François *et al.* 2009;<br>Miranda *et al.* 2010; Gao *et al.* 2018) |
| *HYM1* | 3 | 4 | RAM signaling | no | yes (Nelson *et al.* 2003;<br>Song *et al.* 2008; Saputo *et al.* 2012) |
| *CBK1* | 3 | 4 | RAM signaling | no | yes (Nelson *et al.* 2003;<br>Song *et al.* 2008; Saputo *et al.* 2012) |

\**SIT4* regulates several processes including multidrug resistance and sphingolipid synthesis (Khandelwal *et al.* 2018).

**Figure 2.**
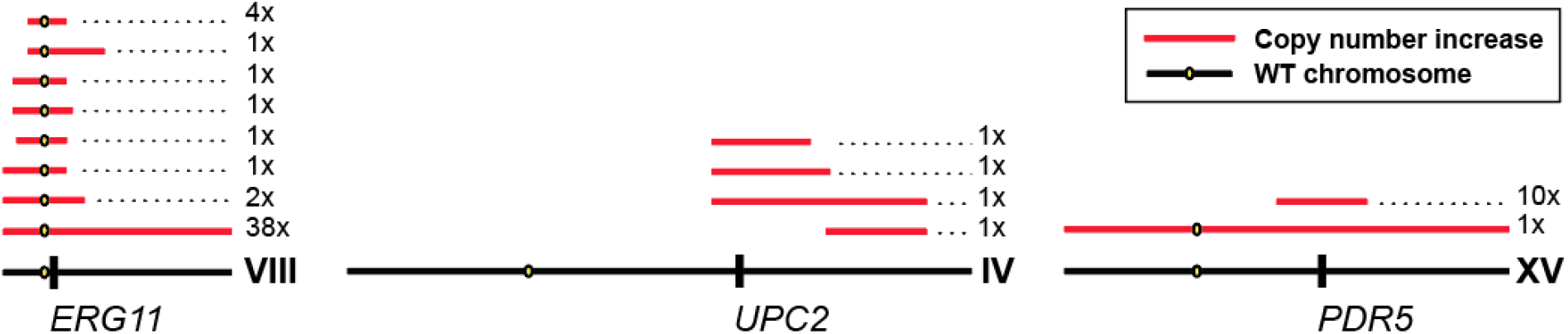
Copy number variation events and candidate genes. Likely segmental duplications on chromosomes VIII, IV, and XV based on increased coverage in whole-genome sequencing data (**Methods**). Regions with increased copy number in at least one sequenced clone are represented with red horizontal lines. Number of clones with a given amplification are listed to the right of each red line. Location of candidate genes that could contribute to a fitness benefit are denoted on wild type chromosome in black.

As a quality check on both the evolution procedure and the sequencing analysis, we first looked in our high frequency mutations for genes known to confer azole resistance (**Table 1**). We found every sequenced clotrimazole-resistant clone had at least one of the following: (1) a nonsynonymous mutation in *PDR1* or *PDR3*, which are likely gain-of-function based on their occurrence in diploids as well as haploids; (2) a loss of the mitochondrial genome leading to a petite phenotype; or (3) a DNA copy number increase involving chromosome VIII, which contains *ERG11*, the target of azoles. The DNA copy number changes included both whole chromosome aneuploidy as well as smaller segmental duplications of chromosome VIII that center around *ERG11* (**Figure 2**; **Table S9**).

We then expanded our analysis to more completely survey the mutations found.

### Copy number variants were common and may impact azole resistance factors

The majority of clones (61 of 99) had at least one copy number variant (CNV). The most common CNVs involved chromosomes I (7 clones), II (4 clones), IV (4 clones), VIII (49 clones), and XV (11 clones) and were most often full chromosome amplifications or segmental duplications (**Figure 2**; **Table S9**). Additionally, two of the five sequenced diploid strains had decreased coverage of chromosome I, indicating a loss of one copy of that chromosome. Aneuploidy has been shown to impact azole resistance in *Candida albicans* due to increased copy number of the genes *ERG11* and *TAC1* (Selmecki *et al.* 2006; Selmecki *et al.* 2008; Selmecki *et al.* 2009). *ERG11* is on *S. cerevisiae* chromosome VIII, the most frequently amplified chromosome across sequenced clones, making this gene an attractive candidate for providing the observed fitness benefit. This is supported by the fact that all of our segmental duplications include *ERG11*. Candidate resistance genes are included within all segmental duplications in (**Figure 2**; **Table S9**).

### Loss of mitochondrial genome was common in evolved clones

The loss of the mitochondrial genome (aka petite or ρ^0^ status) occurred in 73% (72 of 99) of drug-resistant clones. These observations were made based on examination of sequence alignments to the reference mitochondrial genome. Petite mutants have been shown to have an increased resistance to azole drugs (Hallstrom and Moye-Rowley 2000; Ferrari *et al.* 2011), with the tradeoff that they are unable to grow using non-fermentable carbon sources. To confirm that these were petite mutants, all clones were cultured using a medium containing glycerol, a non-fermentable carbon source (**Table S8**). We identified one clone (Westridge_M_final_1_2018-2019) with a wild type mitochondrial genome that did not grow in glycerol medium, indicating it had lost respiratory activity due to a nuclear mutation. This strain possessed unique nonsynonymous mutations in the genes *ERG25*, *TDA9*, and *YKR078W*, as well as a non-coding mutation 276 bases upstream of the gene *SLP1*.

### Transposable element mobilizations biased toward azole resistance factors

We identified only 14 transposable element mobilizations across all 99 sequenced isolates. Only one gene had multiple (four) mobilization events that interrupted its sequence: *SUR1*, an enzyme involved in sphingolipid production which impacts membrane composition, a mechanism by which Sur1 impacts azole resistance (Francois *et al.* 2009). Another mobilization event was detected in the 3’ end of *PDR3*, which encodes a transcription factor regulating the pleiotropic drug response.

### Point mutations in well-characterized azole resistance factors

One goal of the evolution experiments was to isolate clones with multiple mutations that could impact azole resistance. Strains from Moscow High School had an average of 4.8 point mutations, Westridge School early clones had 5, and Westridge School late had 11.6. One strain, from the late timepoint at Westridge School, had a particularly high number of mutations (48, compared to the next highest with 23). Since clones from Westridge School were isolated at multiple time points, some clones shared mutations. We have denoted shared mutations as non-independent events, which can be found in **Table S6**.

In total, 575 unique point mutations were detected across all 99 clotrimazole resistant isolates. Of these, 466 were nonsynonymous, indel, or nonsense mutations. A GO term enrichment analysis on this subset (using the GO term finder tool at https://yeastgenome.org/goTermFinder) found clusters of genes related to drug binding, DNA binding, and regulation of metabolic processes, but with diverse cellular functions (**Table S7**). Even among these nonsynonymous mutations, we anticipate that a nontrivial fraction will be neutral: in other experimental evolution contexts only 20% (Buskirk *et al.* 2017) to 35% (Payen *et al.* 2016) of all mutations found in evolved strains were estimated to be beneficial. To gain further insight into mechanisms of resistance, attention was focused on genes with three or more independent nonsynonymous or nonsense point mutations, since these are more likely to be causative as opposed to passenger mutations, which should be more randomly distributed (**Table 1**). Mutations in this subset of genes accounted for 33.9% of all detected mutations. Many of these genes have well-characterized connections to azole resistance (such as *ERG25* and *PDR5*) which we discuss below and summarize in (**Table 1**).

Half of the azole-resistant clones (49 of 99) possessed missense mutations in *ERG25*. None of the mutations in *ERG25* in the experiment were clear null alleles (nonsense or frameshift). We saw mutations in, and copy number increases centered around, *UPC2* and *PDR5* (**Figure 2**; **Table S9**)*. UPC2* encodes a master regulator of ergosterol synthesis and *PDR5* encodes a major drug efflux pump regulated by Pdr1 and Pdr3. Gain-of-function mutations in both these genes have been shown to impact azole resistance (Flowers *et al.* 2012 (*UPC2*); Uniprot (*PDR5*)).

We also detected mutations in genes that impact sphingolipid production such as *SUR1*, *CSG2*, and *SIT4* (Gulshan and Moye-Rowley 2007; Baudry *et al.* 2001), in addition to the four transposable element mobilizations interrupting *SUR1* (**Table S10**). Sphingolipids interact with ergosterol to produce lipid rafts and perturbation of sphingolipid levels is thought to impact azole accumulation (Francois *et al.* 2009). *SIT4* has been shown to impact expression of the multidrug resistance pathway as well (Miranda *et al.* 2010).

We detected four independent point mutations in *TAO3*, which encodes a component of the conserved RAM kinase signaling network (Nelson *et al.* 2003). In addition, we detected three independent mutations in RAM network genes *HYM1* and *CBK1* and two each in *KIC1* and *SOG2*. To our knowledge none of these genes have a reported azole phenotype in *S. cerevisiae* (SGD, accessed 8/30/2019), but zinc finger transcription factor Ace2, which is regulated by the RAM network (specifically by phosphorylation by Cbk1), has been implicated in both increased azole susceptibility (miconazole) and azole resistance (fluconazole) (Vandenbosch D *et al.* 2013; Kapitzky L *et al.* 2010). In *S. cerevisiae*, null mutations of *ACE2* confer an increased azole resistance when yeast are growth on agar media (Kapitzky *et al.* 2010), and a decreased resistance when grown as biofilms (Vandenboosch *et al.* 2013). Also, deletion of *MOB2* of the RAM network has been shown to cause increased susceptibility to conazoles (Guan *et al.* 2020). Literature from other species of yeast indicates that RAM network component knockouts impact azole resistance in a species or growth phase-dependent manner (Song *et al.* 2008, Saputo *et al.* 2012, Walton *et al.* 2006, Mulhern *et al.* 2006, Homann *et al.* 2009). Several of the mutations from our experiments result in early stop codons (two of four *TAO3*; one of three *HYM1*; zero of two *KIC1*; one of two *SOG2*; zero of three *CBK1*), though the majority of these truncate less than 10% of the encoded protein.

Evolved strains were enriched for mutations in the heme-regulated transcription factor genes *HAP1* and *ROX1* (Zhang and Moye-Rowley 1999). These genes are regulated by oxygen (Kwast *et al.* 1998) and in turn regulate a variety of cellular processes including expression of genes involved in ergosterol synthesis (Serratore *et al.* 2018). The lab strain we utilized for our evolution experiments (S288c) has a transposable element insertion near the 3’ end of *HAP1* that reduces the functionality of the Hap1 protein (Gaisne *et al.* 1999). Nine of the *HAP1* mutations we detected are frameshift and sixteen are nonsense, indicating that further loss-of-function leads to the resistance phenotype. Two of the six detected *ROX1* mutations are an early stop and a single base deletion leading to a frameshift (Y204* and I39indel), suggesting that these are null mutations. Indeed, deletions of *ROX1* have been shown to increase azole resistance (Henry *et al.* 2002).

We detected an enrichment of missense mutations in the genes *ATP1* and *ATP2*, which function in the mitochondrial F1F0 ATP synthase (Saltzgaber-Muller *et al.* 1983; Takeda *et al.* 1986). This complex plays an important role in cellular respiration by synthesizing ATP from the electrochemical gradient generated by the electron transport chain (Nilsson and Nielsen 2016), so it is possible that these mutations impact metabolism in a way that is similar to or synergistic with petite status. Indeed, null alleles of *ATP1* have been shown to exhibit a petite phenotype independent of mitochondrial genotype status (Takeda *et al.* 1986), and all of our *ATP1* and *ATP2* mutants lost their mitochondrial genome. Importantly, mutations in components of the F1F0 ATP synthase, including *ATP1*, have been shown to increase expression of *PDR5* (Zhang and Moye-Rowley 2001).

### Recurrent mutations in mRNA degradation and an uncharacterized mitochondrial protein

In all clotrimazole-resistant isolates, the prevalence of mutations in genes with known roles in drug resistance supported the efficacy of our selection protocols. However, many additional mutations enriched in several genes have not explicitly been implicated in azole resistance. All nonsynonymous mutations identified in this study were enriched for the P-body GO term (**Table S7**). This includes genes that decap and degrade inactive mRNAs in processing bodies (P-bodies) (Sheth 2003; Wickens 2003; Nissan and Parker 2010). Specifically, six mutations were identified in the gene *DHH1* and four in *DCP2*, that encode an activator of mRNA decapping and a decapping enzyme, respectively (Nissan and Parker 2010). Two additional mutations were identified in the 5’-3’ exonuclease *XRN1* that degrades uncapped mRNAs (Larimer and Stevens 1990). The majority of other mutations related to P-bodies had an unclear impact on encoded protein function; two mutations in the catalytic N-terminal domain of *XRN1*^S1155*^ and *XRN1*^C201indel^ are likely loss-of-function or null mutations based on their location in this crucial domain (Larimer and Stevens 1990). Further, deletions of *XRN1* (Kapitzky *et al.* 2010)(Gao *et al.* 2018) and *DHH1* (Vandenbosch *et al.* 2013) have been shown to increase azole resistance in separate genome-wide deletion collection screens.

The only other gene with four or more mutations that does not have a clear connection to azole resistance is *CSF1*. The function of this gene is unknown, though it has been linked to an inability to ferment at low temperature (Tokai *et al.* 2000) and it is a conserved gene from yeast to humans. To test whether *CSF1* mutations impact azole resistance, we introduced one of these mutations (A2913P) into a wild-type strain using a CRISPR/Cas9 genome editing strategy and competed it against a wild type strain and a strain harboring a synonymous mutation in *CSF1* (**Methods; Figure 3A**). We mixed *CSP1*^A2913P^, the synonymous mutant, and the original ancestor in equal proportions and grew these in media with or without 9μM clotrimazole (**Figure 3A**). We found that the *CSF1*^A2913P^ mutation fixed in media containing clotrimazole but not in YPD, indicating that this mutation improves the fitness of *S. cerevisiae* under the selective pressure of an azole antifungal drug (**Figure 3B**).

**Figure 3.**
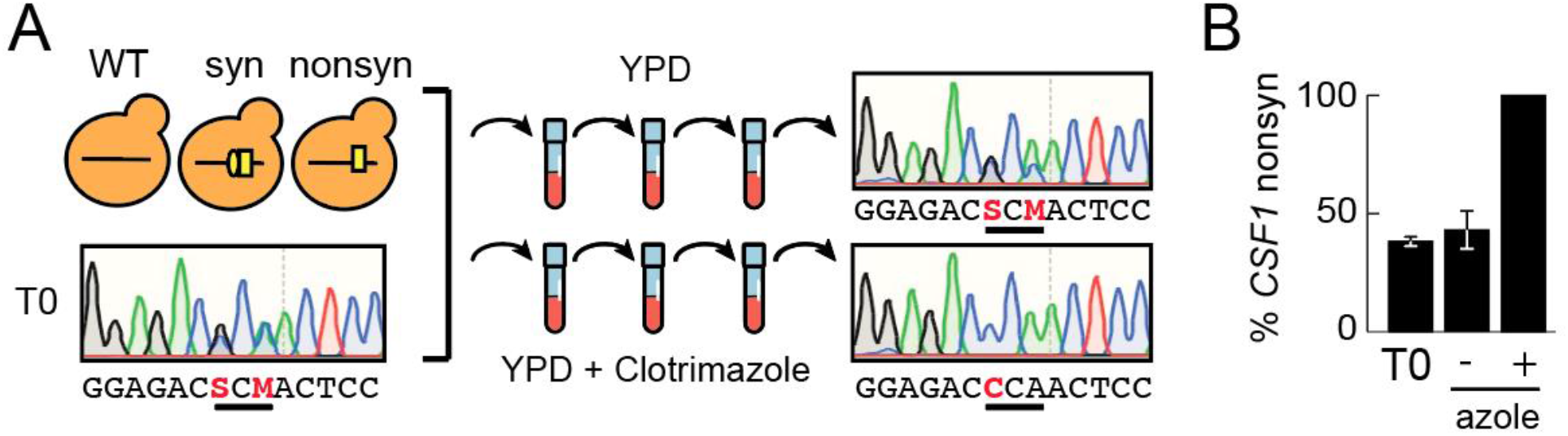
Outline of CSF1 competition experiment. (A) Wild type, synonymous mutant, and nonsynonymous mutant (*CSF1*^A2913P^) were mixed in equal ratios and inoculated into YPD growth media with or without clotrimazole. These populations were propagated for 3 outgrowths. Frequency of *CSF1*^A2913P^ was determined at initial and final timepoints by Sanger sequencing. Representative Sanger sequencing chromatograms are shown. Heterozygous positions are represented with IUPAC codes in sequences below chromatograms (S used when G and C present; M used when A and C present) (B) Frequency of *CSF1*^A2913P^ allele at beginning and end of competition with (+) or without (-) clotrimazole. Frequencies are averages of 3 replicates and were quantified by the program QSVAnalyzer. Error bars are one standard deviation in each direction.

### Evidence of epistatic interactions between adaptive mutations

The large number of replicates available allowed us to look for patterns of mutation exclusion and co-occurrence in our evolved clones. These instances can be due to epistatic interactions, which often indicate a functional connection between genes (Lehner 2011; Costanzo *et al.* 2019 PMID: 30901552).

Almost all (91 of 99) clones possessed a mutation in either *PDR1* or *PDR3*. Despite the prevalence of these mutations and the length of our selection protocol, no evolved clone possessed mutations in both *PDR1* and *PDR3*. This may indicate that once a gain-of-function mutation has occurred in one of these paralogs, there is no benefit (or even a negative consequence) to having a second in the context of these experiments. To test this hypothesis, we designed a crossing scheme to generate recombinant progeny in which mutations in both *PDR1* and *PDR3* segregated. To aid in this effort, we examined our list of mutants for strains with (1) opposite mating types, (2) minimal other mutations, and (3) *PDR1* and *PDR3* mutations that could be genotyped by restriction enzyme digestion. Strains Westridge_T_early_1_2017-2018 (PDR3^T949A^) and Westridge_S_early_1_2017-2018 (PDR1^F749I^ and HBT1^T202I^) fit these characteristics. These strains were mated to form a diploid, sporulated, and 16 tetrads were dissected; all segregants were then genotyped and monitored for growth in the presence of 9μM clotrimazole (**Methods**; **Figure 4**). Strains with mutations in both *PDR1* and *PDR3* showed very similar growth rates to strains with a mutation in only one, suggesting that a second mutation does not increase or decrease fitness in the presence of clotrimazole. This result may explain the absence of double mutants among sequenced clones.

**Figure 4.**
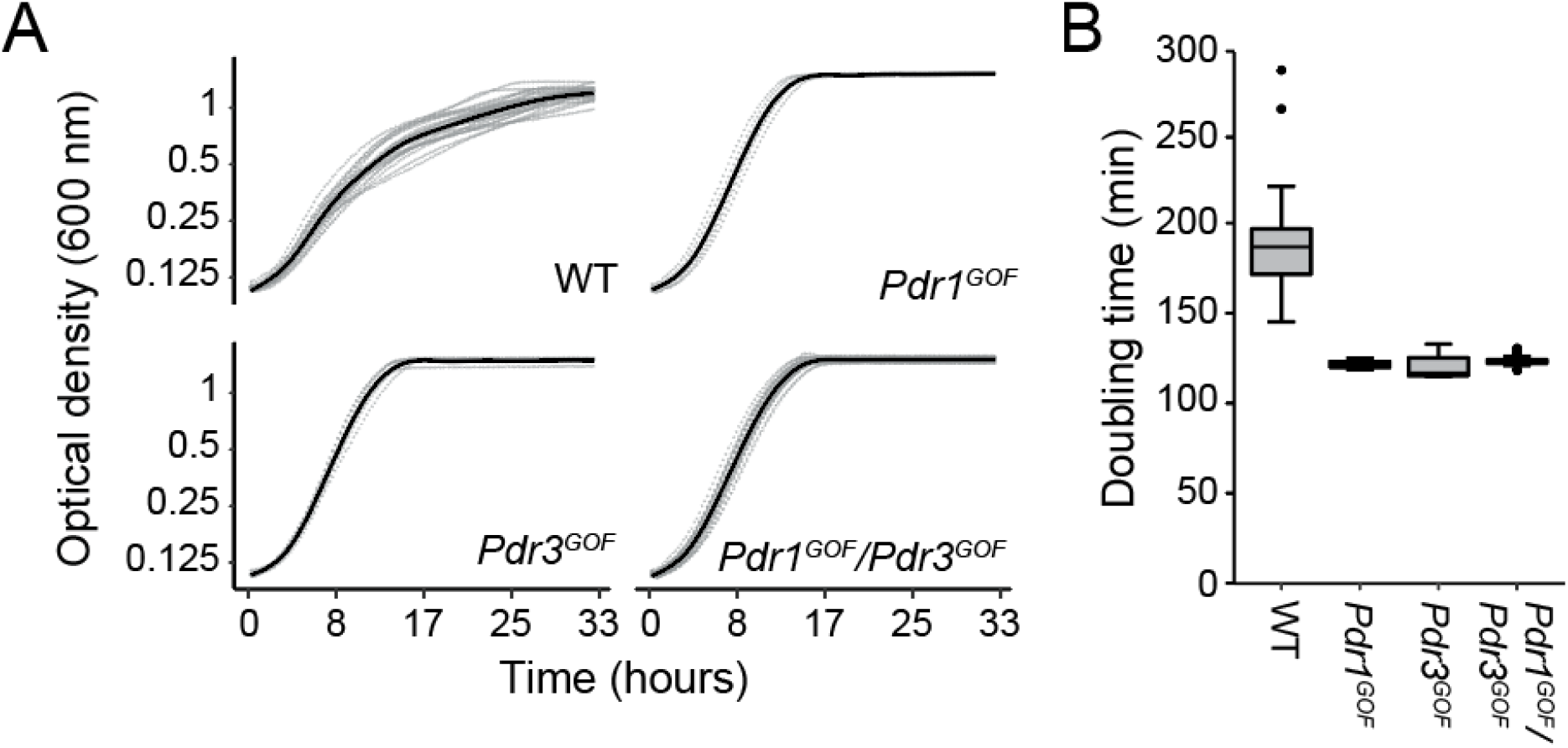
Analysis of growth rates of *PDRX^GOF^* single and double mutants. Haploid *PDR1^GOF^* and *PDR3^GOF^* evolved strains were crossed and sporulated to generate recombinant haploid spores. Spore genotypes were determined by restriction fragment length polymorphisms (**Table S12, Methods**). (A) Segregants were arrayed in a 96-well plate and grown in 9μM clotrimazole media at 30°C in a Biotek Synergy H1 plate reader that measured growth of each strain by optical density (**Table S13**). (B) Average doubling time of each genotype was calculated using the R package *growthrates*.

We also observed that mutations in *HAP1*, *ROX1*, *ATP1*, and *ATP2* co-occurred with mutations in *ERG25* (**Figure 5A**). Specifically, *ERG25* mutants seemed more likely to have secondary mutations in either *HAP1* or *ROX1* and either *ATP1* or *ATP2*. Though approximately half of sequenced clones have a mutation in *ERG25* (49 of 99), only five mutations in *HAP1*, *ROX1* (2x), *ATP1*, and *ATP2* occurred in an *ERG25* wt background, compared to 35 that had a mutation in *ERG25*. In the presence of clotrimazole, *ERG25* mutations generally occurred by early time points, followed by mutations in *HAP1*, *ROX1*, *ATP1*, or *ATP2* which were primarily identified at late time points. This ordering is supported by the higher prevalence of strains with mutations in only *ERG25*, and that clones from lineages with the same *ERG25* mutation have different mutations in *HAP1*, *ROX1*, *ATP1*, or *ATP2* (**Figure 5B**).

**Figure 5.**
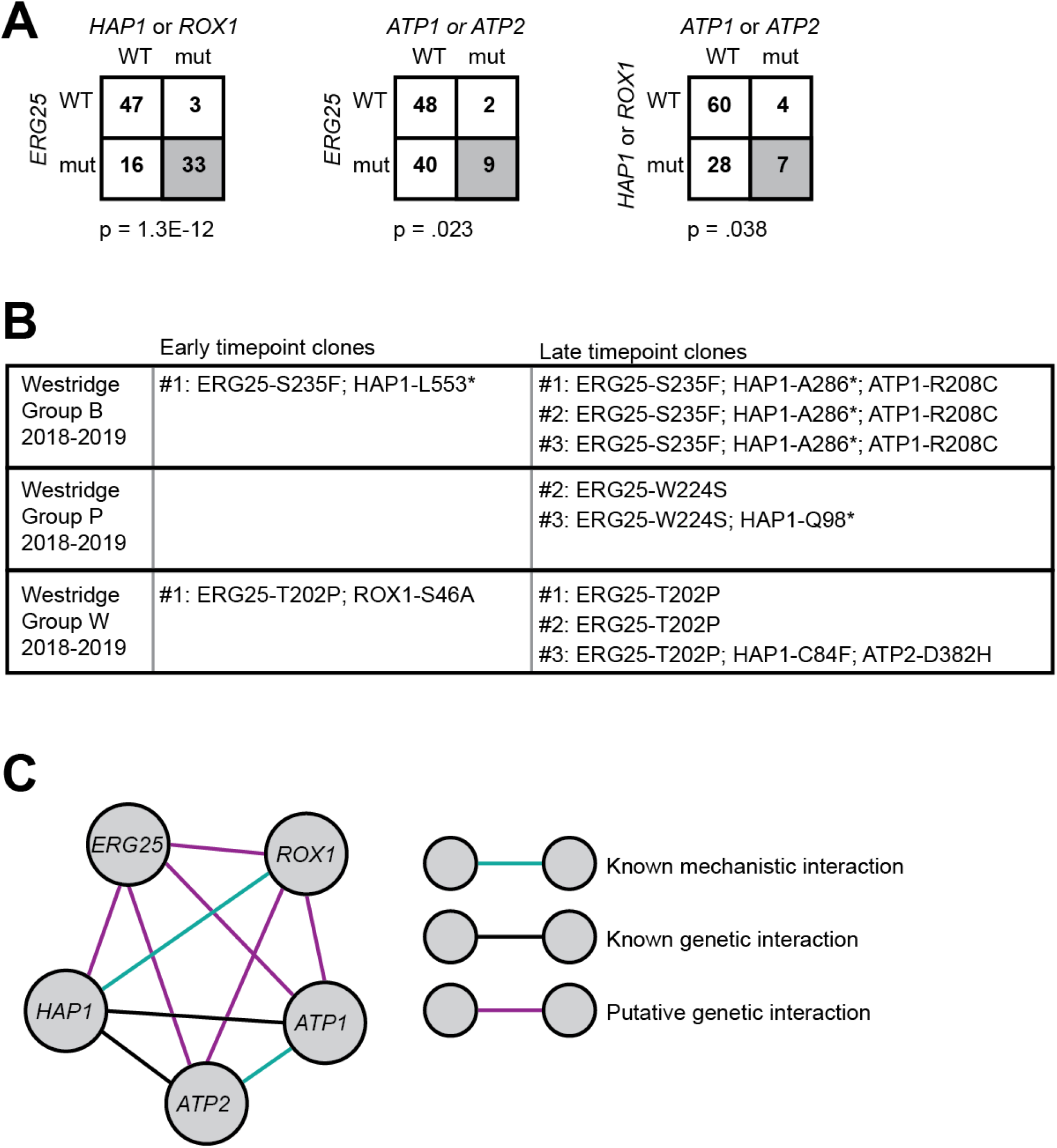
Signatures of epistasis involving *ERG25*. (A) Number of mutants with genotypes related to the genes *ERG25*, *HAP1*, *ROX1*, *ATP1*, and *ATP2*. Frequency is biased toward double mutant groupings in the bottom right of each square (p-values listed below). (B) Lineages within individual replicates were identified by shared mutations in clones from that replicate. Several lineages are made up of clones with *ERG25* mutations that later acquire mutations in *HAP1*, *ROX1*, *ATP1*, or *ATP2*, as evidenced by multiple individuals with the same *ERG25* mutation but different mutations in *HAP1*, *ROX1*, *ATP1*, or *ATP2*. No lineages were detected in which a *HAP1*, *ROX1*, *ATP1*, or *ATP2* mutant later acquired an *ERG25* mutation. (C) Blue lines are known mechanistic relationships (Hap1 regulates Rox1; Atp1 and Atp2 are in the same complex). Black lines are previously-identified genetic interactions (Leeuwen *et al.* 2016). Purple lines are putative novel genetic interactions supported by presented data.

## Discussion

In this paper, we demonstrated how course-based research experiments with high school students can yield insight into how selectable changes at the DNA level lead to changes in molecular factors that impact measurable increases in the organism-level phenotype of azole resistance. In addition to furthering our science research goals, student involvement advanced their understanding of how genotype and phenotype intersect, which is critical to understanding the process of evolution (NRC 2012; NGSS Lead States 2013). We expand on our pedagogical goals and outcomes from this project in (Taylor *et al.* in preparation).

From a basic biology standpoint, long-term evolution experiments provide time for multiple mutations to occur in a single strain, which increases the efficiency of mutant identification by sequencing. These experiments also allow for mutations with smaller or epistatic effects to occur. As evidence of these advantages, all of the novel *CSF1* mutations we identified were found in strains isolated from later time points, as well as much of the evidence implicating many of the processing body mutations and much of the evidence of epistatic interactions. Utilizing alternative genetic backgrounds or altered environments may prove fruitful to expand upon our understanding of azole resistance. Together, our findings underscore the utility of long-term selection to isolate diverse mutations that can impact azole resistance. Increasing the replication of these experiments can improve their power to detect novel adaptive mutations and epistatic interactions, making them ideal for large-scale replication in a classroom setting.

### Concordance with pathogenic isolate sequencing and prior genetic screens

Many of the mutations we identified are in-line with findings from sequencing of drug-resistant pathogenic species of yeast isolated in clinical or agricultural settings (e.g. (Ford *et al.* 2015 E. Life; Paul and Moye-Rowley 2014)). Like the clones from our experiments, drug-resistant fungi frequently possess gain-of-function mutations in orthologs of PDR1 and PDR3, copy number variations that lead to increased copy number of the ERG11 gene, or mutations in regulators of ergosterol production such as UPC2. Copy number variants akin to what we observed in S. cerevisiae have been shown to impact azole resistance in experimental evolution and clinical isolates of the pathogenic yeast Candida albicans (Selmecki *et al.* 2006; Selmecki *et al.* 2008; Selmecki *et al.* 2009). Clinical and environmental isolates of pathogenic fungi frequently have point mutations in ERG11 (Fisher *et al.* 2018 Science) that alter the interaction of the enzyme’s active site with azoles, which we did not detect in our experiments.

Surprisingly, we have not observed mutations in ERG3, which are frequently identified in pathogenic yeasts with azole resistance. Mutations in ERG3 are thought to prevent the accumulation of the fungistatic azole intermediate 14 alpha methylergosta 8-24 (28) dienol, produced as the result of Erg11 inhibition by azoles. Instead, we saw mutations in the enzyme Erg25, which is involved downstream of Erg11 in ergosterol biosynthesis. Mutations in Erg25 may act to decrease activity of the gene in some way, as decreased expression has been shown to increase azole resistance (Smith *et al.* 2016). Similar to Erg3, loss of Erg25 function has been shown to suppress toxicity associated with null mutations in Erg11 or inhibition of Erg11 by azoles. Loss of Erg11 and Erg25 function is thought to enable the incorporation of lanosterol as an alternative to ergosterol in the fungal membrane. Importantly, it was shown that reduced heme biosynthesis suppressed this toxicity (Gachotte *et al.* 1997) and is consistent with the appearance of non-functional alleles of HAP1 and ROX1 that encode regulators of heme biosynthesis.

Petite mutants of *Candida glabrata* have been isolated in clinical settings (Bouchara *et al.* 2000) and may have increased *in vivo* virulence (Ferrari *et al.* 2011). These mutants are unable to undergo cellular respiration due to mutations that impact mitochondrial function, which has been shown to increase the activity of Pdr1 and Pdr3 through an unspecified post-translational mechanism (Traven *et al.* 2001; possible mechanisms in Shahi *et al.* 2007 and Shahi *et al.* 2010). These mutants are thus unable to grow on non-fermentable carbon sources. Rho^0^ (loss of mitochondrial genome) petite mutants were common across the clones we sequenced, with 72 of 99 clones having experienced this type of mutation. We identified one clone which displayed a petite phenotype (inability to grow in glycerol-containing media; a non-fermentable carbon source) despite possessing its wild type mitochondrial genome, suggesting that its phenotype is due to a nuclear mutation. This strain possessed unique nonsynonymous mutations in the genes *ERG25*, *TDA9*, and *YKR078W*, as well as a non-coding mutation 276 bases upstream of the gene *SLP1*. *TDA9* is a transcription factor that regulates acetate production, and *TDA9* null mutants produce less acetic acid during wine fermentation (Walkey *et al.* 2012). *TDA9* has a paralog, *RSF2*, which encodes a gene that regulates growth in glycerol-containing medium (Lu *et al.* 2005). These details make it an attractive candidate for causing the observed petite phenotype.

### Connection between P-bodies and azole resistance

Though P-bodies have not explicitly been implicated in azole resistance, two of our candidate genes (*XRN1* and *DHH1*) have been implicated in separate genome-wide deletion collection screens, and a genome-wide mutagenesis study (Gao *et al.* 2013) found an enrichment for the P-body GO term, which was undiscussed in their paper. It is unclear how these mutations would impact azole resistance at a mechanistic level, though intriguing candidates exist. Evidence suggests that gain-of-function mutations in *CaCDR1* (*PDR1/PDR3* ortholog) impact the stability of its transcripts (Manoharlal *et al.* 2010; Khakhina *et al.* 2018), and that mitochondrial activity regulates these genes through a post-transcriptional mechanism (Travern *et al.* 2000; Shahi *et al.* 2007; Shahi *et al.* 2010). Perturbation of mitochondrial function and application of clotrimazole alters the frequency and morphology of P-bodies (Buchan *et al.* 2011). Loss-of-function mutations in P-body components such as Dhh1, Dcp2, and Xrn1 may then prevent degradation of *PDR1* and *PDR3* transcripts. Such a mechanism would likely have pleiotropic effects on other traits, since P-bodies accumulate many mRNAs.

### Connection between CSF1 and azole resistance

We observed recurrent mutations in *CSF1*, which encodes a protein of unknown function that localizes to the mitochondria (Dubreuil *et al.* 2019) (Reinders *et al.* 2006). Mutations in *CSF1* impact fermentation specifically at low temperatures (Tokai *et al.* 2000). In a genome-wide screen *CSF1* was implicated in maturation of glycosylphosphatidylinositol (GPI), a post-translational modification that allows proteins to be targeted to the cell membrane (Čopič *et al.* 2009). Genes involved in this maturation process have been shown to regulate *ERG11* in *Candida albicans* and can thus modulate azole resistance (Jain *et al.* 2019). Genome-wide genetic interaction mapping experiments have shown that the interaction profile of *CSF1* is similar to that of many genes involved in cell surface GPI anchor maturation (**Table S14**)(Usaj *et al.* 2017). Moreover, *csf1Δ* has defects in cell wall glycans and is sensitive to membrane and cell wall damaging agents as well as the K1 killer toxin, which is also suggestive of a role in cell wall/membrane integrity (Pagé *et al.* 2003). Further, deletions of *CSF1* have been shown to suppress the azole-resistance phenotype of *erg3Δ* through an unspecified mechanism (Mount 2018). *CSF1* is conserved across many higher eukaryotes. The human gene is hypothesized to be involved in endocytic recycling (Kane *et al.* 2019), and endosomal trafficking mutants have been related to azole resistance in candida (Peters *et al.* 2017). The human homolog is also associated with neonatal death and developmental delay (Kumar *et al.* 2020), so clarifying its mechanism of action and role in cellular function will be of interest.

### Large number of replicates reveals epistatic interactions

We also found novel combinations of mutations that indicate epistatic interactions between resistance mutations. Epistatic interactions tend to reflect an underlying mechanistic connection between involved genes (Lehner 2011; Costanzo *et al.* 2019). These interactions can thus provide insight into the basic molecular biology of networks of genes involved in drug resistance phenotypes. Knowledge of epistatic interactions can allow researchers to predict genes that are essential for resistance evolution (Lukačišinová *et al.* 2020).

Elucidating the mechanism of the interaction between mutations in Pdr1 and Pdr3 may provide interesting new insights into azole resistance. Single gain-of-function mutations or overexpression of each of these genes individually leads to a dramatic increase in expression of the drug efflux pump *PDR5*, among other resistance factors. It is thus surprising that a second mutation does not provide some additional benefit. This may be due to a tradeoff that overrides the beneficial impact of the second mutation. Pdr1 and Pdr3 are paralogs (products of an ancient gene duplication) that regulate an overlapping but distinct set of genes related to multidrug resistance and iron metabolism (DeRisi *et al.* 2000; Tuttle *et al.* 2003). They are additionally regulated in overlapping but distinct ways: for instance, loss of mitochondrial genome impacts *PDR3* but not *PDR1* (Hallstrom and Moye-Rowley 2000; Zhang and Moye-Rowley 2001). Many pathogenic yeasts have only a single copy of the ancestral version of these transcription factors, but many of the regulatory associations with these genes are conserved (Khakhina *et al.* 2018). Clarifying the mechanism of this epistatic interaction may thus provide insight into resistance mechanisms in pathogenic species. It may additionally provide insight into the forces that shaped the evolution of these paralogs after the ancestral gene was duplicated.

The co-occurrence of mutations in the genes *ERG25, HAP1, ROX1, ATP1, and ATP2* suggests that these mutations also interact in some way. Based on their frequency and evidence from lineages with identical *ERG25* mutations (**Figure 5B**), it is likely that in most cases *ERG25* mutations occur first, followed by mutations in *HAP1* or *ROX1*, and finally *ATP1* or *ATP2*. It is thus possible that *ERG25* mutations have a stronger effect on their own, or that mutations in these other genes have a stronger effect when in an *ERG25* background. Hap1 regulates Rox1 expression (Keng 1992), and they can act as activators or repressors of *ERG* gene expression under different conditions (Serratore *et al.* 2018). *HAP1* mutations have previously been shown to co-occur with mutations in *ATP1* and *ATP2,* and have been speculated to be suppressors of a fitness defect from perturbation of the F1F0 ATP synthase (Leeuwen *et al.* 2016). Together, these observations support that these patterns of co-occurrence are indeed due to epistatic interactions representing an underlying mechanistic connection (**Figure 5C**). The mutants isolated from these experiments will be of value for investigating the mechanisms of these interactions.

### Effectiveness of yEvo as a scalable course-based research experience

Recent decreases in the cost of whole-genome sequencing have made evolve-and-resequence paradigms more accessible in a broader range of settings. Our classroom protocols demonstrate the power and robustness of this approach in high school classrooms, which could also be applicable to undergraduate courses (Govindan et al. 2020). These experiments provide a compelling demonstration of the process of evolution in a pedagogical setting, since they can be explored at many levels of biological organization. Other research areas may also benefit from this type of a student-teacher-scientist partnership (Houseal *et al.* 2014) in advancing both pedagogical and biomedical research goals.

## Acknowledgements

We thank the following students for conducting evolution experiments and literature searches: Katie Bender, Maya Bluthenthal, Penelope Boone, Ciauna Cota, Nicole Gibbs, Siena Giljum, Maddie Groff, Io Jette-Kouri, Emily McLane, Makana Meyer, Haley Pak, Alex Perez, Sophia Ramirez-Brown, Lara Sachdeva, Sofia Santoro, Nicole Tanouye, Sarah Arellanes, Chloe Daniel, Sofia Flores-Rojas, Corah Forrester, Summer Garrison, Kathryn Huang, Sophia Kaplan, Helena Karafilis-Spensley, Caroline Nowak, Zellie Owen, Caroline Rygg, Catherine Su, Ashley Wang, Audrey Wang, Sylvia Woolner, Nadina Wu, Lauren Baydaline, Jessica Beskind, Deijah Bradley, Sophia Bulander, Kate Crowell, Ryo Goodman, Rachel Harris, Lucy King, Hannah Lam, Olivia Molina-Kong, Molly Mulane, Kallie Papanikolas, Dalia Rizkana, Andrew Lee, Isabella Lee, Frances Fletcher, Christine Panahi, Bella Gilchrist, Ava Horner, Mackenzie Bowlen, Jaya Sadda, Abigail Yuhan, Anelise Pardo, Solaar KirkDacker, Sophie Carter, Christine Balian, Jasmin Palomo, Zaynab Eltaib, Juliane Zanides, Sydney Flashman, Leah Soldner, Sophia Lew, Rosalie Lansing, Hattie Bilson, Keira Myles, Ayiana Saunders-Newton, Elisa Dong, Nicole Aivazis, Abbey Piatt Price, Monika Lee, Vivian Lee, Amanda Tse, N’Dea Piliavin-Godwin, Kathleen Honey, Zahra Vogel, Erisa Rosen, Mia Hakian, Olivia Bulow, Hailey Lam, Hana Odawara, Isabella Welling, Cara Wilson, Lily Yu, Claire Denault, Claire Coffman, Tate Ahn, Natalie Darquea, Maribella Munoz-Jimenez, Jennifer Spinoglio, Eva Bassel, Emily Hsieh, Caris Lee, Zelia Mauro, Shirlynn Chan, Bayley Dickinson, Jacqueline Yipp, Haley Ansel, Monica Lopez Ramos, Camilla Connor, Emerson Thein, Krystal Raymundo, Julia Cruz, Coco Goran, Audrey Ma, Natalie Chen, Mable Zhang, Liv Bjorgum, Eleanor Washburn, Alina Chiu, Isabella Grigorian, Amin Rezamand, Jana Veleva, Serena Strawn, Jaston McClure, Eliza O’Murphy, Sophie Gomulkiewicz, Hannah Jenkins-Evans, Linnea Sheneman, Emilia Fountain, Elle Rasmussen, Ellie Gomulkiewicz, Aengus Kennedy, Noah Gregg, Kira Vierling, Laurel Hicke, Ammon Kunzler, Evan Odberg, Samantha Hammes, Ava Jakich-Kunze, Kendall Forseth, Christina Petrie, and Eric Thorsteinson. We additionally thank Renee Geck for extensive edits on the manuscript; Jef Boeke and Jasmine Temple for color plasmids; Westridge School and Moscow High for participating; Ken Berger and Lee Anne Ereckson for protocol development and experiment supervision; Dianne Newman and Melanie Spero for assistance making yeast media; Sayeh Gorjifard for figure design assistance; Atina Cote and Fritz Roth and their students Bean Raktan Ahmed, Avery Albert, Amanda Black, Bryan Bombon Moreno, Rachel Bradley, Ouye Chen, Michelle Cheung, Natasha Dhamrait, Annette Diao, Emily Durant, Lauren Durland, Isabella Gallello, Linsey Gong, Mila Gorchkova, Emily Hoover, Sornnujah Kathirgamanathan, Jack Daiyang Li, Tamara Li, Jongmin Lim, Daniel Martinho, Jacob McAuley, Winona McGregor, Ksenia Meteleva, Janice Mwangi, Sabina Pang, Bruno Pereira, Sakshi Shinghai, Michelle Tello Calle, Matthew Tran, Jhenifer Vasquez Rojas, Ran Xu, Justin You, Shilei Zeng, Qing Fang Zhang for sequencing a subset of the clones presented in this manuscript.

## Funding

This work was supported by National Science Foundation grant 1817816. This material is based in part upon work supported by the National Science Foundation under Cooperative Agreement No. DBI-0939454. Any opinions, findings, and conclusions or recommendations expressed in this material are those of the author(s) and do not necessarily reflect the views of the National Science Foundation. The research of MJD was supported in part by a Faculty Scholar grant from the Howard Hughes Medical Institute. MBT and CRLL were supported by T32 HG000035 from the National Human Genome Research Institute.

## Conflicts of Interest

The authors declare no conflicts of interest.

## Notes

### Competing Interest Statement

The authors have declared no competing interest.

